# Bayesian inference of infectious disease transmission from whole genome sequence data

**DOI:** 10.1101/001388

**Authors:** Xavier Didelot, Jennifer Gardy, Caroline Colijn

**Affiliations:** Department of Infectious Disease Epidemiology, Imperial College London, Norfolk Place, London, W2 1PG, United Kingdom; Communicable Disease Prevention and Control Services, British Columbia Centre for Disease Control, Vancouver, British Columbia, Canada; School of Population and Public Health, University of British Columbia, Vancouver, British Columbia, Canada; Department of Mathematics, Imperial College, London SW7 2AZ, UK

## Abstract

Genomics is increasingly being used to investigate disease outbreaks, but an important question remains unanswered – how well do genomic data capture known transmission events, particularly for pathogens with long carriage periods or large within-host population sizes? Here we present a novel Bayesian approach to reconstruct densely-sampled outbreaks from genomic data whilst considering within-host diversity. We infer a time-labelled phylogeny using BEAST, then infer a transmission network via a Monte-Carlo Markov Chain. We find that under a realistic model of within-host evolution, reconstructions of simulated outbreaks contain substantial uncertainty even when genomic data reflect a high substitution rate. Reconstruction of a real-world tuberculosis outbreak displayed similar uncertainty, although the correct source case and several clusters of epidemiologically linked cases were identified. We conclude that genomics cannot wholly replace traditional epidemiology, but that Bayesian reconstructions derived from sequence data may form a useful starting point for a genomic epidemiology investigation.

---

Infectious disease outbreak investigation typically involves both field work – interviews that establish links between patients and/or exposures to a source of infection – and molecular epidemiology, in which laboratory typing of pathogen isolates is used to identify related cases. Data from both streams are considered together to reconstruct the outbreak, identifying its origins and pathways of onward transmission. Ideally, the reconstruction aids in the real-time management of the outbreak and guides public health policy and practice in preventing future occurrences.

Molecular epidemiology has recently undergone a revolution with the advent of next-generation genome sequencing. With the cost of sequencing a complete bacterial genome now comparable to that of a gold-standard typing analysis [1], traditional molecular methods for investigating bacterial disease outbreaks are increasingly being replaced by genomics [2–5]. Whereas techniques such as multilocus sequencing typing interrogate approximately 0.1% of a bacterial genome, whole-genome sequencing permits identification of sequence variation across the complete genome. Given the short timescales over which outbreaks typically occur, only a small number of single nucleotide changes are expected between outbreak isolates. This diversity is not captured by traditional typing methods, but can be identified and leveraged for outbreak reconstruction using whole-genome approaches.

Despite the emergence of genomic epidemiology, reconstructing outbreaks is not straightforward. Most reconstructions published to date rely heavily upon fieldwork data, and the utility of genome sequencing for inferring transmission events in the absence of these data – which, for a given outbreak, may be incomplete or unavailable – remains unclear, particularly for diseases characterized by periods of latency or chronic infection, during which substantial within-host genetic diversity may arise. Although most investigations include a phylogenetic analysis of the outbreak isolates, phylogenetic trees do not directly correspond to transmission trees and cannot, as such, identify specific person-to-person transmission events [6]. In a densely-sampled outbreak, all cases will appear as tips of the phylogenetic tree, when in actuality some of these cases will have infected others, so internal branching events are also associated with sampled hosts.

Nevertheless, it should be possible to infer transmission from sequence data using alternative approaches, and several such methods have been proposed [7–13]. These methods typically identify transmission events as branching events in a phylogeny, and they do not consider within-host genetic diversity. This simplification may be appropriate for pathogens with a fairly small generation time – the time that elapses between a host becoming infected and causing other infections – but not for those with long generation times, latency, and carriage. In these organisms, one can expect several nucleotide changes to accrue within a single host, with different individual lineages being transmitted onwards to secondary cases. Examples include *Clostridium difficile* [14, 15], *Mycobacterium tuberculosis* [16, 17], *Staphylococcus aureus* [18–20] and *Helicobacter pylori* [21, 22]. Excluding the possibility that multiple distinct genetic lineages of a pathogen – distinguished by only a few mutations – can be present within a host can lead to serious misinterpretation of putative transmission events [23].

Furthermore, current methods for transmission inference do not leverage the capabilities of time-calibrated phylogenetic inference methods. Given the importance of factors such as branch length in inferring the underlying host contact network structure from a phylogeny [24], an inference method that incorporates sampling times and a molecular clock analysis is preferable to one using neighbour-joining, maximum parsimony, or other simplified tree-building algorithms. With the increasing use of genomic epidemiology, a method capable of inferring transmission events from genomic data alone whilst also considering within-host diversity offers an important tool for outbreak reconstruction. Here, we present a Bayesian inference scheme based on timed phylogenetic trees that can be run independently of epidemiological data or weighted with whatever fieldwork data is available, such as timing and duration of infectivity, level of infectivity, and geographic location. Our method rests on a within-host pathogen population genetic model, under which different lineages may be transmitted to secondary cases. We first reconstruct a timed phylogenetic tree and then infer the underlying transmission network, given the observed phylogeny. This two-step approach represents a formalisation of a previous approach to reconstruct transmission events on top of a timed phylogeny [25]. We make use of existing robust methods for phylogenetic inference such as BEAST [26] and ClonalFrame [27], and employ a novel Bayesian methodology to infer a transmission network via a Monte-Carlo Markov Chain (MCMC). We apply our method to simulated data to illustrate its performance, and to a real-world dataset to reconstruct the transmission of *Mycobacterium tuberculosis* in a densely-sampled outbreak.

## Results

### Within-host diversity affects placement of transmission events on a phylogenetic tree

To assess the impact of within-host diversity on the inference of transmission events from a phylogeny, we simulated a transmission tree – the set of specific person-to-person transmission events within an outbreak – and genealogies arising from this tree under two scenarios of effective population size. We modelled an outbreak in a susceptible population of size *N* = 100, with a per-year recovery rate *γ* = 2 and per-year, per-contact infectivity rate *β* = 0.02. The expected number of infected individuals in a SIR model is *R*(*∞*), such that 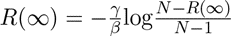 [28], which here is *R*(*∞*) = 13.51. A transmission tree *T* was simulated under these conditions, with 10 individuals infected (Figure 2A).

**Figure 1.**
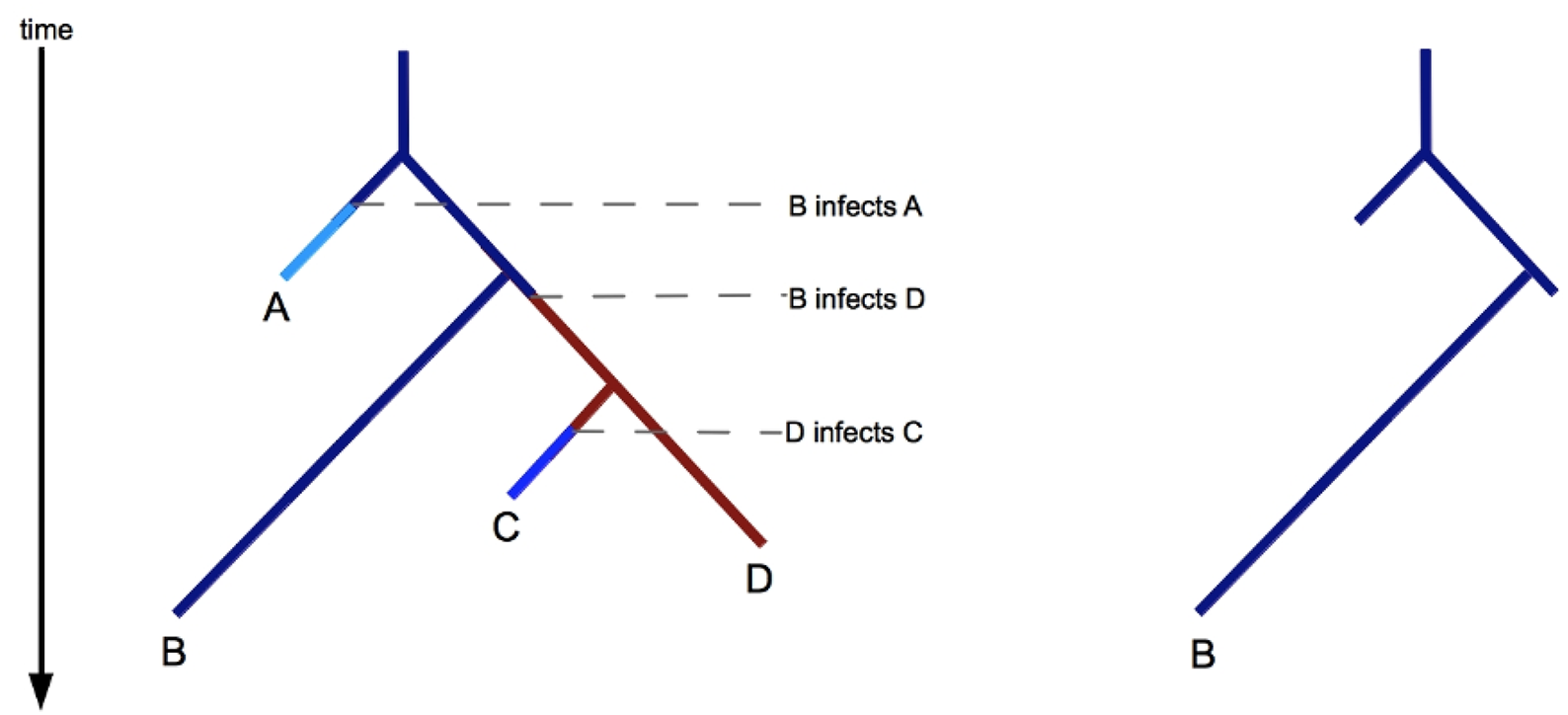
The coloured genealogical tree (left). Each host isolate corresponds to a unique colour. A lineage is coloured according to the host it was in at the corresponding time. When a lineage changes from colour *c_i_* to *c_j_* (forward in time), this represents *i* infecting *j*. Each colour may not persist in the tree after the time of the corresponding tip, because this is the recovery time of the host. The sub-tree restricted to a single colour (right) is the part of the tree inside the corresponding host; lineages are taken from this tree at the recovery time and the times when the host infected others.

**Figure 2.**
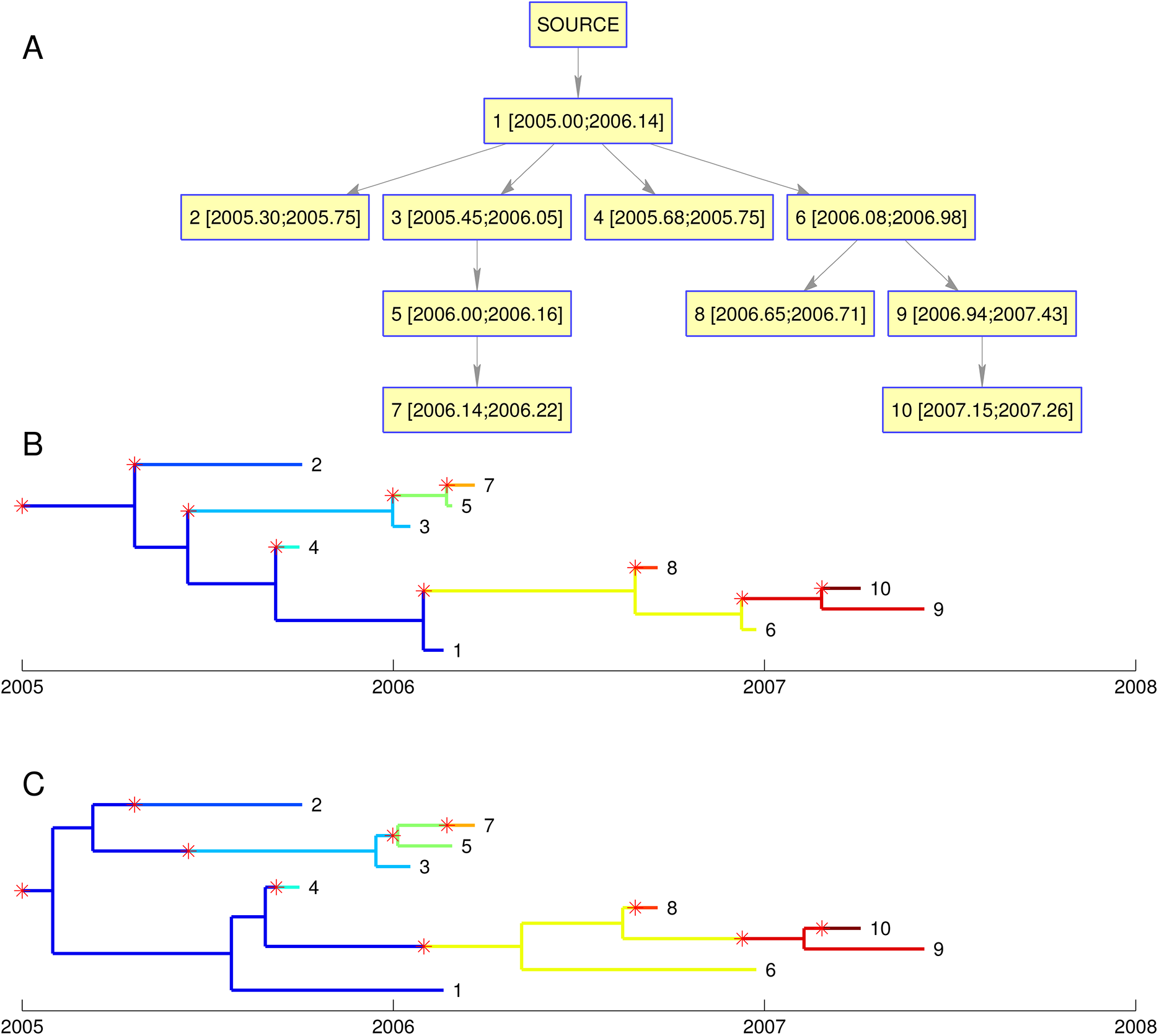
(A) Transmission tree simulated with *N* = 100, *γ* = 2 and *β* = 0.02. The numbers in square brackets represent respectively the time of infection and removal of each individual. (B) Genealogical tree simulated conditional on the transmission tree in part A and with parameter *N_e_g* = 0. Transmission events are indicated by red stars and a change in branch colour. (C) Same as in part B but with parameter *N_e_g* = 0.274.

If the effective population size within hosts is *N_e_* = 0, then within-host diversity is zero and transmission events coincide exactly with coalescent events of the phylogeny (Figure 2B). This assumption simplifies the relationship between the transmission tree *T* and the genealogy *G* [13] and may be appropriate for infections with a short incubation time. However, in cases of latent disease, chronic infection, or long carriage periods, this assumption may not be valid. An example of this is asymptomatic carriage of *Staphylococcus aureus*, which can persist within a host for months to years [29–31] so that the recovery rate *γ* = 2 above is realistic, but the parameter *N_e_g* (effective population size *N_e_* times duration of a generation *g*) is on the order of 100 days [18, 19].

Simulating a genealogy *G* under the same transmission tree *T* but with *N_e_g* = 100*/*365 = 0.274 year illustrates the complex relationship between *G* and *T* that arises under a model of within-host diversity (Figure 2C), with transmission events no longer corresponding to coalescent events. In particular, it is now possible for the genomes from two hosts A and B to share a common ancestor more recently than with the genome from a third host C, even if C infected both A and B. An example of this in Figure 2C is provided by host 1, who infected both hosts 2 and 3. In spite of this, 2 and 3’s genomes are more closely related to each other than to 1’s, because both 2 and 3 were infected by a lineage from 1 that is different from the one that was sampled. If no within-host diversity was assumed, the genealogy in Figure 2C precludes the possibility of 1 infecting both 2 and 3. Under our more realistic model with *N_e_g* = 0.274, these transmissions become a possibility.

### MCMC inference of a transmission network from a phylogeny captures known transmission parameters and events

To determine whether the known events and parameters of transmission tree *T* could be inferred from a genealogy *G*, we applied our MCMC method to the simulated dataset in Figure 2C. The MCMC was run for 100,000 iterations with the first half (burn-in) discarded. This took approximately 5 minutes on a desktop computer, and several independent runs were compared to ensure good convergence and mixing of the chain.

The inferred rate of infectivity *β* had posterior mean 0.021 (95%CI 0.010-0.038) capturing the correct value of *β* = 0.02, while the inferred rate of removal *γ* had posterior mean 1.82 (95%CI 0.88-3.17), also capturing the correct value of *γ* = 2. The inferred effective population size *N_e_g* had posterior mean 0.49 (95%CI 0.02-2.49), which included the correct value of *N_e_g* = 0.274, but that was also compatible with *N_e_g* values up to an order of magnitude higher or lower than the correct value. This result reflects the difficulty of precisely inferring *N_e_g*, especially as only ten individuals were infected. In a separate simulation with *β* equal to 0.05, such that *R*(*∞*) = 89 hosts became infected, all parameters were more precisely reconstructed, with CIs for *β*, *γ* and *N_e_g* of 1.9-2.4, 0.04-0.06, and 0.19-2.11, respectively.

The posterior sample of transmission trees inferred based on the phylogeny in Figure 2C was summarised using a graph representing all transmission events with posterior probability above 10% (Figure 3A). All ten correct transmission events are present, but incorrect events are also contained in the graph, particularly the reverse of correct edges, making it difficult to distinguish infector from infected. With 30% of the posterior probability, host 1 was correctly identified the most likely source case (Figure 2A), but hosts 2 and 3 were also likely sources, with 25% and 17% respectively. Overall, 33% of the posterior probability weight was carried by correct edges, with the remaining two-thirds of probability supporting transmission events that did not actually occur in the simulation. We used Edmonds’ algorithm [32] to find the spanning tree within the graph carrying the maximum posterior weight to identify the most likely transmission scenario. In the resulting point estimate of the transmission tree, seven of the ten simulated transmission events were recovered (Figure 3B).

**Figure 3.**
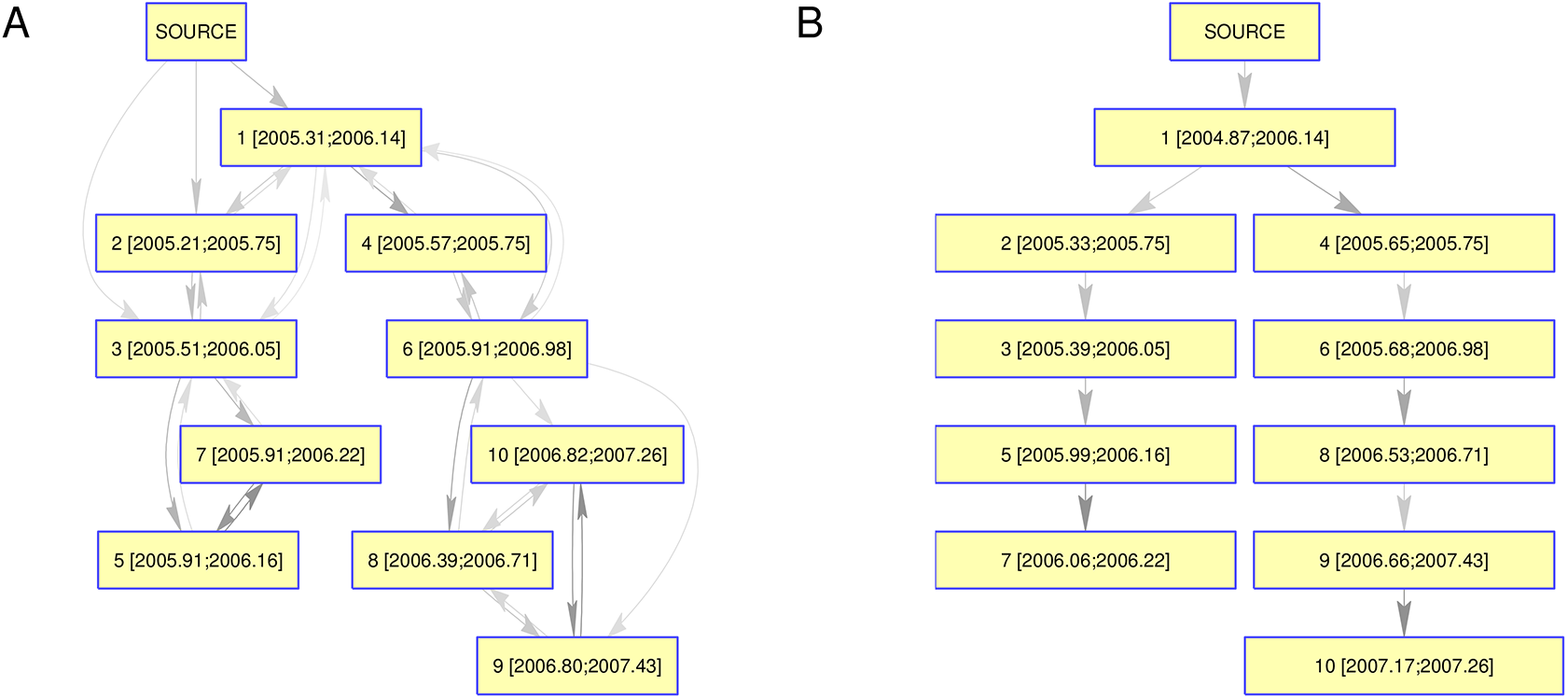
(A) Network representation of the posterior distribution of transmission trees for the simulated dataset shown in Figure 2C. Nodes represent hosts, with in square brackets to mean inferred infection time and the known removal time. Edges represent a posterior probability of transmission of at least 10%, with a darker edge indicating a higher probability. (B) Point estimate of the transmission tree based obtained by taking the maximum spanning tree in the network shown in part A.

One hundred scenarios analogous to the one shown in Figure 2 were generated, each using the same parameters *β*, *γ* and *N_e_g* and each including ten infected individuals. Inference was performed for each of the hundred simulated datasets. On average, we found that correct transmission events carried 27% of the posterior probability weight. In each simulation, there are only ten correct transmission events (including transmission from an external source to the first case out of a total of a hundred possible events (each of the ten hosts can infect any of the other nine, plus ten choices for transmission from the external source to the first case). Correct edges therefore carry, on average, almost four times more weight than incorrect edges ((0.27/10)/(0.73/100) = 3.7).

### Reconstruction of a *Mycobacterium tuberculosis* outbreak dataset

To assess the performance of our inference method on a real-world dataset, we applied it to an outbreak of tuberculosis for which a hypothesized source case had been identified and for which epidemiological data supporting several further transmission events was available. A BEAST phylogeny (Figure 4A, Figure S1) indicated that most of the inferred transmission events occurred after the source case (K02) recovered, meaning the source could have infected eight other people at most – eight being the number of unique lineages present in the tree at the source’s recovery time – and that the source harboured significant within-host diversity. The clock rate estimated by BEAST had a posterior mean of 1.15*×*10*^−^*^7^ (95%CI 0.39*×*10*^−^*^7^ to 2.00*×*10*^−^*^7^), consistent with previous estimates in *M. tuberculosis* [17, 33].

**Figure 4.**
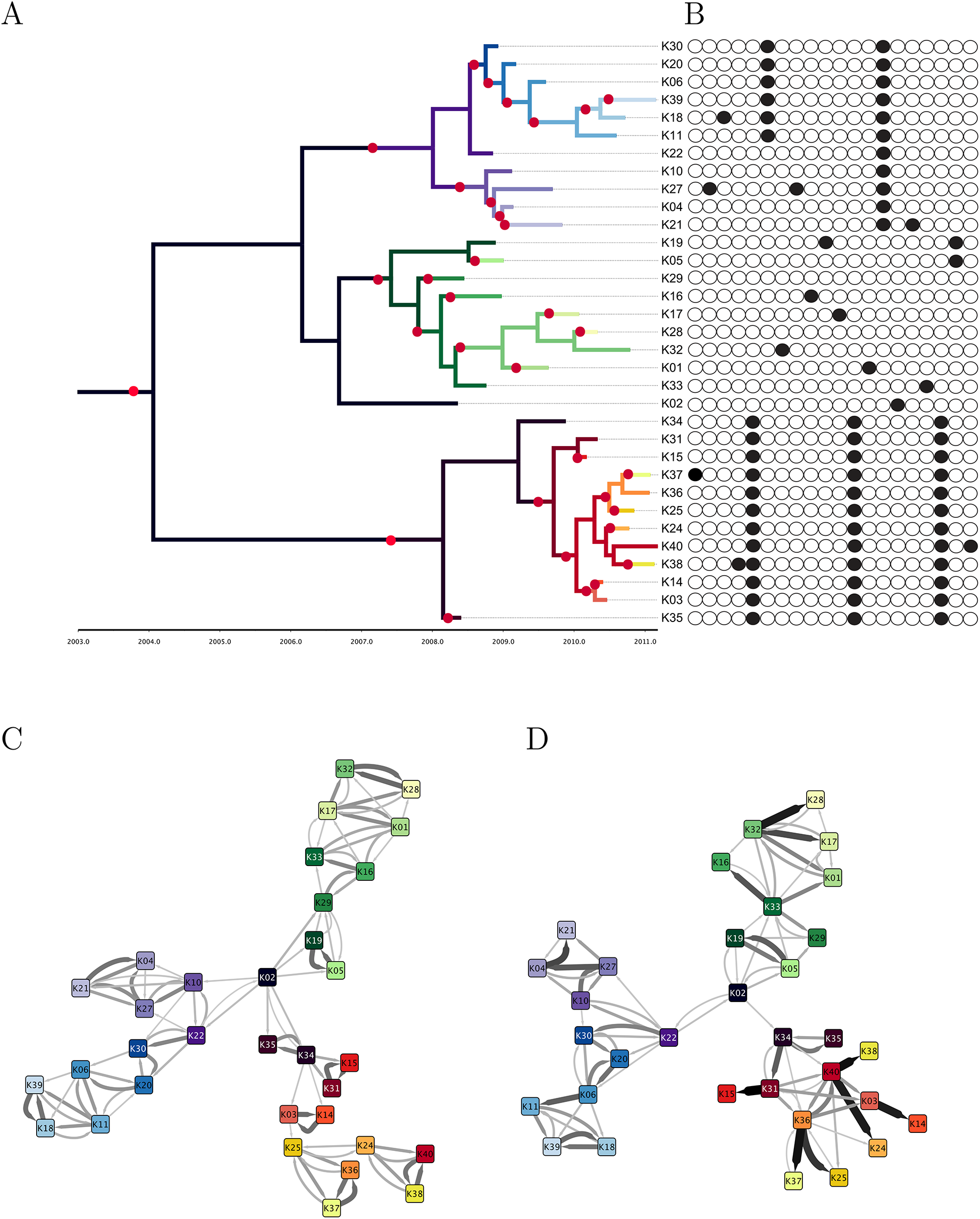
Application to a real-world tuberculosis outbreak. (A) Phylogenetic tree inferred by BEAST. (B) SNVs differentiating the isolates. (C) Transmission network inferred without epidemiological modification. (D) Transmission network inferred with epidemiological modification. In parts (C) and (D), edges are shown with width and shading proportional to their posterior probability, except edges with low probability which are omitted. In part (A), the phylogenetic tree is coloured according to the consensus transmission tree from part (D) and using the same unique colours for each each host as in parts (C) and (D).

We used the MCMC approach to infer transmission networks under two scenarios: one based only on the phylogenetic tree (Figure 4C), and one in which posterior probability weights were modified using epidemiological data (Figure 4D), including infectivity, infectious period, and geographic location. While both graphs placed the suspected source K02 as central to the network, the modified network displayed both a lower clustering coefficient (0.483 versus 0.596) and a higher average per-edge posterior probability (0.302 versus 0.253), meaning that the modified network contained fewer bidirectional events, displayed a more web-like layout suggestive of waves of transmission rather than chains, and returned more high-probability events.

Comparing inferred transmission events to known epidemiology revealed that both the basic and modified networks did capture aspects of known epidemiology, but in different ways. Amongst eight contacts sharing a sleeping space with the source case, three were identified by both reconstructions, two were present only in the basic reconstruction, and one was present only in the modified reconstruction. Another reported household contact between two cases was not returned in the basic reconstruction, but was present in the modified version. Both inferred networks recapitulated a cluster of three cases linked to a specific sleeping location; in the basic model the cluster appears as a complete subgraph with roughly equal probabilities for all six edges, while in the modified version it is clear that a single individual infected two other cases. We also examined the 15 transmission events with the highest probabilities in the modified reconstruction and found that three were well-supported by epidemiological information, seven were possible given the known locations and contacts of cases, two did not appear to have an epidemiological linkage between the predicted infector/infected, and three were highly unlikely.

The three unlikely transmissions involved the three cases known to reside in a different geographic location than the other outbreak cases and thought to have been infected by a visiting traveller from the outbreak community. In the basic reconstruction, this scenario is mostly recapitulated – the traveller infects two further cases. In the modified version, despite including an adjustment for geographic location, the traveller does not infect any of the other three cases, nor are they infected by individuals with a travel history to the other community.

The parameters estimated with and without epidemiological modification were similar, with (*β*, *γ*) = (0.0037, 1.29) and (0.0032, 1.23) respectively. This value of the removal rate *γ* indicates that, on average, 15 months elapsed between infection of a host and its detection. We interpret the transmission rate *β* in terms of the basic reproduction number *R*_0_ = *βN/γ*, the expected number of secondary infections caused by a case in a susceptible population. Here, we calculate *R*_0_ = 1.15 (95%CI 0.73-1.9) in the basic reconstruction and *R*_0_ = 1.03 (95%CI 0.65-1.69) in the modified reconstruction. This relatively low number is expected for tuberculosis in a low-burden, highly resourced setting like the one studied here [34].

For simplicity, the networks described above were based on a single phylogeny, namely the maximum clade credibility tree returned by BEAST. However, there was significant uncertainty in the BEAST posterior, as illustrated by a DensiTree plot [35] (Figure S2). We therefore inferred transmission trees separately for each of one hundred trees sampled by BEAST, and the aggregated results are shown in Figures S3 and S4 for the application without and with epidemiological modification, respectively. In the case of the latter, accounting for the uncertainty in the phylogenetic tree does not result in a great increase in uncertainty for the reconstructed transmission tree.

## Discussion

We present a Bayesian inference method for reconstructing transmission events in a densely sampled outbreak using time-labelled genomic data. By modelling within-host evolution as a neutral coalescent process, we are able to account for within-host genetic diversity and accurately ascribe infection events to a host harbouring multiple lineages of a pathogen – an issue critical to the reconstruction of diseases with latent or asymptomatic carriage periods and/or chronic infection. We first reconstruct a phylogeny, leveraging the strengths of existing packages for phylogenetic inference such as BEAST [26], and next infer a transmission tree using an efficient MCMC procedure which can be run on a posterior sample of phylogenies and hence incorporate phylogenetic uncertainty.

Recently, an approach was proposed to relate phylogenetic trees to transmission trees [23], using a similar decomposition to ours; they simultaneously inferred a phylogeny and a transmission tree for a dataset on 12 farms infected with foot-and-mouth disease [11]. Their focus is on sampling and joint inference given a specific within-host population model and a sharply defined incubation period. In contrast, our method is very rapid and can be applied to large (densely sampled) outbreaks, it is applicable to infections with long and variable infectious periods, it exploits the capabilities of BEAST, and infers rather than specifies the in-host effective population size. We are also able to flexibly incorporate additional field epidemiological data of various types.

Our results demonstrate that even if the genomic data is perfectly informative about the phylogeny, as in our simulations, considerable uncertainty is present in the transmission network when a realistic model of within-host evolution is taken into account. While true events are captured, they are often eclipsed by other potential events and cannot be completely identified, even with methods that extract the subgraph with maximal posterior probability weights. This contrasts with previous approaches to outbreak reconstruction which equated phylogenetic branching with transmission [11, 13] and may be too assertive when within-host diversity exists. Our Bayesian approach captures this uncertainty, and permits the integration of genomic data with epidemiological field data; any epidemiological information can be integrated into the prior for the transmission tree without affecting the remainder of the method.

When applied to a real-world dataset, the method correctly inferred the most likely source case and several key transmission clusters supported by field data. Neither the basic nor the epidemiologically-modified networks recapitulated all suspected transmission events, and the modified version in particular failed to return several strongly suspected transmissions. This emphasizes the important point that genome sequencing in a public health context will not make epidemiological data redundant, but instead should be seen as a complementary stream of information. When applying the method to actual outbreak data, running the inference under both basic and modified scenarios is recommended, followed by careful scrutiny in the light of available field data. However, it is important to note that overall patterns of transmission are of significant interest to public health. Our method suggests that genomic information can shed light on whether outbreaks tend to occur as long chains of transmission or as large bursts from single hosts, and whether transmissions tend to be associated with specific environments or hosts, even if individual transmission events remain uncertain.

Like any model-based statistical analysis, our method makes a number of assumptions, some of which might not be appropriate for every application. The within-host evolutionary dynamic is modelled by a simple coalescent process with constant population size [36]. This could easily be relaxed to allow for variation in the within-host population size following infection whilst staying in a coalescent framework [37]. However, our specification represents a natural choice since it is the simplest possible choice of model with only a single parameter *N_e_g*. Very little is known about the within-host evolutionary dynamics of most infectious diseases, but the use of a more complex model could be justified for example if it led to a better marginal likelihood as measured by a Bayes Factor [38]. Our model also assumes that the transmission bottleneck is complete. This would be more difficult to relax as any other choice would imply that some genetic diversity rather than a single variant can be transmitted from infector to infected, and therefore that the common ancestor of two isolates from the same host might be in a different host. Again, very little is known about this property for most infectious disease, but amongst the few microorganisms where this was formally investigated, data suggests the bottleneck is very strong if not complete, for example in HIV [39–42].

The main limitation of the method as presented here is that it assumes that all cases comprising an outbreak have been sampled. This assumption was acceptable for the tuberculosis application, given the surveillance and reporting systems in place for TB, but clearly would not be appropriate in all settings – for example, diseases with milder symptoms where cases may go unrecognized. However, our two-step approach provides a good starting point to investigate these outbreaks too. The phylogeny inferred in the first step does not assume full sampling and the colouring in the second step could be redefined so that a colour corresponds not just to a single host, but rather to a host plus all intermediate cases up to the next sampled case. Our model would need to be modified to account for this and additional parameters must be introduced, such as the proportion of unsampled infected hosts, but the general approach of colouring the phylogeny should remain valid, and future work will seek to implement this generalised model.

## Methods

### Model for the epidemiological process between hosts

For the epidemic between-host spreading process, we consider a stochastic, continuous time Markov chain (CTMC) version of the general SIR epidemic model [28]. In a population of known size *N*, the parameters of this process are the infection rate *β* and removal rate *γ*. If the state of the process at time *t* is (*S_t_, I_t_, R_t_*), denoting the number of susceptible, infected and removed individuals, respectively, then transition to the state (*S_t_−*1, *I_t_* + 1, *R_t_*) happens at rate *βS_t_I_t_* and transition to the state (*S_t_*, *I_t_* − 1, *R_t_* + 1) happens at rate *γI_t_*. The former event is a transmission and the latter a removal. The process is started in the state (*S*_0_ = *N −* 1, *I*_0_ = 1, *R*_0_ = 0) and is run until the epidemic finishes at *I_t_* = 0.

This basic process has been extensively studied, especially in a Bayesian framework where inference can be performed using a Monte-Carlo Markov Chain (MCMC) [43, 44]. Let *n* denote the number of individuals who have been infected, ie the number of removed individuals at the end of the process. Let *T* denote the transmission tree, where each node is one of the *n* infected individuals and edges represent transmission events. In our notation, *T* contains the information of who infected whom and when, as well as when individuals were removed. Simulation of a transmission tree *T* given the parameters *N*, *β* and *γ* can be done using the Gillespie algorithm [45].

### Model for the genealogical process within hosts

We consider that when each individual is removed, a single genome from the infectious agent is isolated and genotyped. The genealogical relationships between these genomes can be represented as a timed genealogy *G*, where leaves correspond to the *n* genomes (one from each host), and internal nodes represent common ancestors of the genomes. The genealogy *G* depends on the transmission tree *T*, but also depends on the within-host evolutionary dynamics [46] and the transmission bottleneck [47, 48]. For simplicity, we model within-host evolution as a neutral coalescent process [36] with constant population size *N_e_* and average generation length *g*. In this model, any two lineages within a host coalesce at constant rate *N_e_g*, which is therefore the sole within-host parameter of interest. Furthermore, the transmission bottleneck is assumed to be complete, so that only a single genomic variant is transmitted from infector to infected. This transmitted genome is a random sample from the infector’s pathogen population at the time of transmission.

Under these assumptions, the genealogy *G* can be simulated given a transmission tree *T* and parameter *N_e_g* as follows. In a first step, *n* subtrees are generated corresponding to the genealogical process within each host, and in a second step, these subtrees are pasted together in order to produce *G*. For the first step, the within-host genealogy of each host *i* is simulated independently of each other. The genealogy within host *i* has a single root at the time when *i* was infected, a leaf at the time when *i* was removed, plus a leaf for each host that *i* infected at the time when these transmission events happened. Each within-host genealogy is generated using the coalescent with temporally offset leaves [49] with coalescence rate *N_e_g* and using rejection sampling on the coalescent tree [50] to ensure that a single lineage exists at the time when *i* became infected. For the second step, the pasting is done for each transmission event. For example, if A infected B at time *t* in the transmission tree *T*, then the subtree of A contains a leaf at time *t* and the subtree of B has its root at time *t*, and pasting is done between this leaf and this root. Repeating this pasting for all transmission events completes the generation of the genealogy *G*.

### Bayesian decomposition

Our approach assumes that all *n* infected individuals are known and that their removal times are also known. Based on the genomes sampled from each of the *n* individuals, the timed genealogy *G* is reconstructed, and for the time being we assume that *G* is known exactly. Since the times of the leaves of *G* represent the removal times of the *n* individuals, our notation *G* includes these removal times, but not the times of infection. Due to the within-host genealogical process, transmission events do not occur at branching points in the genealogy. The aim of the inference is to infer the epidemiological parameters *β* and *γ*, the within-host evolutionary parameter *N_e_g* and the transmission tree *T*. The posterior distribution of interest is therefore:

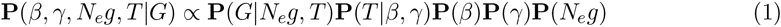

The last three terms represent the prior distribution of the parameters *β*, *γ* and *N_e_g* which we take to be Exponential(1), and it remains to specify **P**(*G|N_e_g*, *T*) and **P**(*T|β, γ*).

First, the term **P**(*G|N_e_g, T)* in Equation 1 represents the probability of the observed genealogy given the transmission tree and within-host parameter *N_e_g*. We relate a transmission tree *T* to the genealogy *G* by associating each point on *G* to a host. This can be visualised as a “colouring” of the branches in *G*: each host *i* is represented with a unique colour *c_i_* and a branch segment is coloured with *c_i_* if it corresponds to evolution that happened within host *i.* Transmission events therefore correspond to point on *G* where the colour changes and do not, in general, coincide with branching times in the genealogy. The same host may harbour several lineages ancestral to our sample at the same time, which can be visualised on the tree. Figure 1 illustrates this visualisation and our notation. For a given host *i*, we can consider the tree *G_i_* obtained by looking only at the branches of *G* that are coloured with *c_i_*. This tree *G_i_* corresponds to the evolutionary process within host *i*, and has a number of leaves *n_i_* equal to one plus the number of hosts infected by *i*.

To find **P**(*G|N_e_g, T)* we exploit the fact that the subtrees *G_i_* correspond to evolution within each host *i* = 1*..n*, and so are independent of each other. Since the distribution is conditional on *T*, the dates *d_i,j_* of the *j* = 1*..n_i_* leaves in *G_i_* are known (corresponding to all transmissions from *i* plus the removal of *i*). The date *r_i_* of the root of *G_i_* is also known since it corresponds to the infection of host *i*. This leads to the decomposition:

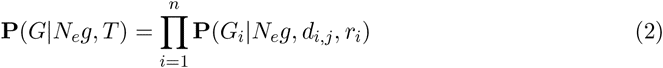

The term **P**(*G_i_|N_e_g, d_i,j_, r_i_*) is the probability under the coalescent model with rate *N_e_g* of the timed genealogy *G_i_* given the dates of its leaves, *d_i,j_*, and conditional on having only one ancestor by the time *r_i_* of the infection of host *i*. If this last condition was not present (ie if *r_i_* was a very long time ago), then this distribution would be that of the coalescent with temporally offset leaves as described in [49]. The additional condition corresponds to the fact that the most recent common ancestor (MRCA) of the leaves has to be more recent than the date of infection since the transmission bottleneck is absolute.

To calculate **P**(*G_i_|N_e_g, d_i,j_, r_i_*), we consider the *n_i_* leaves in increasing order of age. For each leaf *j*, we consider adding it on the genealogy formed by the previous leaves 1*..j −* 1. The first (most recent) leaf corresponds to a linear genealogy with probability 1. The second leaf has to coalesce with the linear genealogy of the first leaf before the infection time *r_i_*. The third leaf has to coalesce with the genealogy formed by the first two leaves before the infection time *r_i_*, etc. This process is repeated for all leaves, so that the genealogy formed at the end is the complete *G_i_*. When adding leaf *j*, let *A_j_* denote the sum of branch lengths in the genealogy formed so far between the time of leaf *j* and the time where it coalesces with an ancestor of a previously considered leaf. Similarly, let *B_j_* denote the sum of branch lengths between the time of leaf *j* and the time *r_i_*. *A_j_* is exponentially distributed with parameter *N_e_g*, but this distribution is truncated by *A_j_ < B_j_* to ensure that coalescence happens before time *r_i_*. We therefore deduce that **P**(*G_i_|N_e_g, d_i,j_*_=1_*_..ni_, r_i_*) can be calculated as follows:

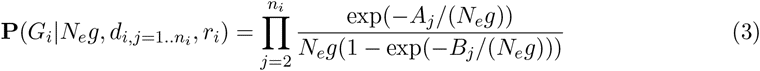

This now completely specifies **P**(*G|N_e_g, T)* from Equation 1.

The term **P**(*T|β, γ*) in Equation 1 represents the probability of the full epidemiological process given the epidemiological parameters. *T* includes the times of all infection and removal times. Let *t_i_*_=1_*_.._*_2_*_n_* represent the list of all these event times (ie *n* infections plus *n* removals), sorted chronologically, and let *e_i_*_=1_*_.._*_2_*_n_* be equal to 0 for infections and 1 for removals. At any time *t*, the number of susceptible, infected and removed individuals is given by:

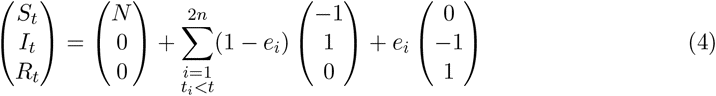

By considering the probability of each event in turn, we deduce that:

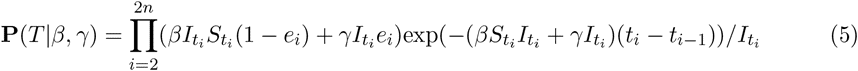

The denominator in Equation 5 comes from the fact that *T* contains not only the states (*S_t_, I_t_, R_t_*) but also knowledge of who was the infector when an infection occurred and who was removed when a removal occurred, for both of which there is a uniform choice amongst the *I_ti_* infected individuals at time *t_i_*.

We now have all the components of Equation 1. Inference proceeds using an MCMC approach.

### MCMC moves

For the parameters of the epidemiological process *β* and *γ*, we use the Gibbs moves introduced by [43]. These moves rely on the fact that the distribution **P**(*T|β, γ*) in Equation 5 has for conjugate prior a Gamma distribution, so that the posterior of these parameters is also a Gamma distribution.

For the parameter of the within-host evolutionary process *N_e_g*, we use a Metropolis move such that the new value is equal to the old one plus a draw from Uniform([*-∈, ∈*]). This move is accepted with probability equal to the ratio of Equation 2 after and before the move, times the ratio of prior probability of *N_e_g* after and before the move.

To update the transmission tree *T*, a single infection event is first chosen at random. If the infection is to the source case, then its date is proposed to be modified by a draw from Uniform([*-∈, ∈*]) (excluding values that would make the transmission to the source case become more recent than the common ancestor of the genealogy *G*). Otherwise, if the transmission is not to the first case, then it corresponds to a transmission event, ie a point at which two colours meet on the genealogy. The proposed MCMC update consists of moving this transmission event uniformly at random to another point on the genealogy where it gives a valid colouring, ie one where (i) there are only *n* − 1 colour changes, (ii) each leaf is coloured in the colour *c_i_* of the host it corresponds to and (iii) the colour *c_i_* does not exist in the tree after the leaf corresponding to host *i*. We note that this move may impact the meaning not only of the transmission event being moved but also of other transmission events on the tree so that the validity of the whole configuration has to be checked. This proposed move is symmetric, and it is accepted with probability equal to the ratio of the products of Equations 2 and 5 after and before the move. In the Supplementary Information we discuss the symmetry in more detail and show that the resulting Markov chain is irreducible.

### Application of the method to a real-world *Mycobacterium tuberculosis* outbreak dataset

Whole genomes of 33 *Mycobacterium tuberculosis* were sequenced on an Illumina HiSeq and short reads were aligned against the *M. tuberculosis* CDC1551 reference sequence using BWA [51]. SAMtools [52] mpileup identified 20 high-confidence single nucleotide variants (SNV) positions that differed amongst the isolates, defined as positions called with a quality score of 222, genotype quality of 99, and no indication of strand basis or low depth of coverage. Phylogenetic inference was performed with BEAST [26] using the concatenated SNV data labelled with sampling dates. A coalescent model with constant population size [36] was used for the tree prior. More complex priors such as a coalescent with exponential growth prior [37] or a birth-death epidemiological prior [53] did not make a significant difference to the resulting phylogeny.

Inference under the basic scenario used Equation 5 for the prior on the transmission tree with *N* = 400, reflective of the number of individuals in the outbreak community estimated to be at risk of tuberculosis exposure. The second scenario modified Equation 5 to incorporate known aspects of the outbreak’s epidemiology, including: the geographic location of a host, host smear status (positive or negative), and whether the transmission event in a proposed transmission network happened at a time that a host was known to be non-infectious (for example, at a time before a negative tuberculosis skin test). We modelled infection at a rate (*β*^+^*I*^+^ + *β^−^I^−^*)*S* and recovery at a total rate *γ*(*I*^+^ + *I^−^*), for which *I*^+^ and *I^−^* are the numbers of smear-positive and -negative cases. We penalised transmission trees in which cases transmit before they are thought to have become infectious based on their clinical history, trees in which cases receive infection before the time of their most recent negative tuberculosis skin test, and trees in which transmission occurs between cases not known to have been in the same location.

This results in modifications to Equation 5, although the principle is the same:

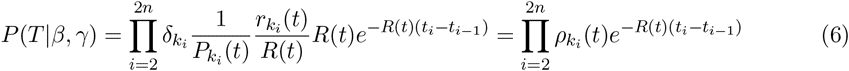

where *R*(*t*) is the total rate at which events occur, so that 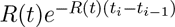 is the probability that time *t_i_* − *t*_*i*−1_ elapsed between events; 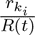 is the probability of the event being type *k_i_* (for example, given that an event occurred after a time interval *t_i_* − *t*_*i* −1_, the probability that it was a recovery is 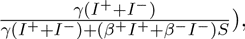 and the population to which the event applies with uniform probability is 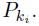 For recoveries, 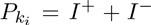 and for infection events it is either *I*^+^ or *I^−^*corresponding to whether a smear-positive or -negative case was the infector. The penalty 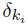 for the event is either 1 (no penalty), 0.05 if the event contradicts the timing of cases’ clinical histories, or 0.005 for a transmission event that occurred when cases were in different locations with no known travel of either case.

The resulting transmission networks for both scenarios were visualized using Cytoscape v.3.0.2 [54], using a yFiles organic layout and visually encoding the posterior probability of an edge using edge colouring and width.

## Acknowledgments

We thank Dr. Rob Parker, Dr. Sue Pollock, Dr. Paul Hasselback, Ms. Lori Hiscoe, and Ms. Denise McKay for their contributions to the outbreak field investigation, and Dr. Patrick Tang, Dr. James Johnston, and Dr. Mabel Rodrigues for their contributions to the genomics work.

